# Lamina and Heterochromatin Direct Chromosome Organisation in Senescence and Progeria

**DOI:** 10.1101/468561

**Authors:** Michael Chiang, Davide Michieletto, Chris A. Brackley, Nattaphong Rattanavirotkul, Hisham Mohammed, Davide Marenduzzo, Tamir Chandra

## Abstract

Lamina-associated domains (LADs) cover a large part of the human genome and are thought to play a major role in shaping the nuclear architectural landscape. Here, we use simulations based on concepts from polymer physics to dissect the roles played by heterochromatin- and lamina-mediated interactions in nuclear organisation. Our model explains the conventional organisation of heterochromatin and euchromatin in growing cells, as well as the pathological organisation found in oncogene-induced senescence and progeria. We show that the experimentally observed changes in the locality of contacts in senescent and progeroid cells can be explained naturally as arising due to phase transitions in the system. Our model predicts that LADs are highly stochastic, and that, once established, the senescent phenotype should be metastable even if lamina-mediated interactions were reinstated. Overall, our simulations uncover a universal physical mechanism that can regulate heterochromatin segregation and LAD formation in a wide range of mammalian nuclei.

## Introduction

The spatial organisation of interphase chromosomes in metazoans is characterised by folding into a hierarchy of structures, from “topologically associated domains” (TADs) [1] to compartments [2] and chromosome territories [3]. TADs are currently understood as originating from the action of processive [4] or diffusing [5] cohesin complexes, whereas the establishment of segregated active and inactive genomic compartments is naturally explained by the polymer-polymer phase separation of chromatin segments bearing similar epigenetic marks [6–10]. At the scale of chromosomes, their territorial nature may be explained by the slow dynamics of chromatin during interphase [11, 12]. Yet, the large-scale nuclear organisation displays a further level of segregation which is less well understood: one in which heterochromatin (HC) is preferentially found in specific concentric layers either near the nuclear lamina (NL) or the nucleoli [13–15], while euchromatin (EC) is enriched in the interior, or middle, layer [14]. Regions of the genome that are preferentially bound to the NL, so-called lamina-associated domains (LADs), are strongly enriched in long interspersed nuclear elements (LINEs) and are associated with gene repression [13]. Intriguingly, LADs display a substantial overlap with nucleolus-associated chromatin domains (NADs) [16], and together they cover more than a third of the human genome.

A common approach to analysing lamina-mediated nuclear organisation is through perturbation studies in which the concentric layering of HC and EC is disrupted. An important example, on which we focus here, is that of cellular senescence. A popular way to trigger senescence is to expose cells to stress – e.g., mitotic stresses or DNA damage. In this way, the cell cycle can be permanently arrested within days [17]. Such cells typically harbour large HC bodies known as senescence associated heterochromatin foci (SAHF) [17, 18]. This nuclear phenotype, which is a hallmark of stress-induced senescence, can be visualised by DNA stains. It is also possible to study cellular senescence by isolating cells from prematurely ageing (progeroid) patients, which harbour a mutation in the lamin A/C (*LMNA*) gene. In both stress-induced senescence and progeria, there is a weakening of the nuclear lamina and of lamina-chromatin interactions; however, qualitative differences between the two states have been reported for markers of HC. In progeria (e.g., in Hutchinson-Gilford Progeria Syndrome, HGPS), there is a reduction in the HC mark histone 3 lysine 9 tri-methylation (H3K9me3), whereas stress-induced senescence seems to be associated with an increase in some HC associated proteins, such as HP1 and Core histone macro-H2A (mH2A), but not H3K9me3. Moreover, progeroid cells are devoid of the SAHF found in stress-induced senescence.

Although SAHF were first identified almost 15 years ago, the connection between the changes in HC proteins and SAHF formation has not been resolved, and we still have no clear understanding of their function, with contradicting suggestions ranging from pro-proliferative activity to irreversible seals of the senescence arrest. In an effort to gain insight into the role of these two key players, lamina and HC, in the nuclear dynamics in cellular senescence, we developed and studied a new model based on concepts from polymer physics. Our model focused on HC-mediated and NL-mediated interactions with chromatin. We generated chromatin mass spectrometry data for senescence to obtain a more quantitative understanding of the changes in HC markers. We analysed chromatin immunoprecipitation with sequencing (ChIP-seq) and LAD DamID data from ENCODE [19] to accurately capture chromosome-NL interactions in human cells. Surprisingly, by varying only two parameters – the strength of HC-HC and HC-NL interactions – our simulations predicted a range of distinct nuclear architectures that are in remarkable qualitative agreement with the known large-scale genome organisation in growing, senescent and progeroid cells. This agreement was then further quantitatively reinforced by high-resolution HiC data, which show that the frequency of long-range interactions markedly increases in senescent cells but decreases in progeroid cells [20].

Having shown that our model can recapitulate the variation in chromatin folding in different cell states by tuning only two parameters, we employed it to shed light on experimentally challenging questions. In particular, our simulations predicted that LADs are stochastic and display cell-to-cell heterogeneity; this provides a natural explanation for the fact that LADs overlap with NADs across a population of cells but are distinct at the single-cell level. In addition, our simulations were able to dissect the dynamics of SAHF establishment upon entering senescence from the growing state. This follows a co-operative process known as polymer desorption [21]. Finally, our model predicted that the growing-senescence transition should be abrupt (as it is thermodynamically first-order-like), so that the senescent state should be metastable even when NL-chromatin interactions are partially re-established. This suggests a biophysical reason for the observation that senescent cells with SAHF do not re-enter the cell cycle.

Our results demonstrate that polymer physics principles can explain the concentric organisation of HC and EC in growing, senescent and progeroid human cells. At the same time, because our model is developed from first principles – i.e. no data fitting from chromosomal contact maps – we can readily export it to study the organisation in other cell types or other organisms. For instance, the inverted organisation found in the nuclei of rod cells of nocturnal mammals [14, 22] entails the desorption of LADs from the lamina and the formation of a dense HC core. It is likely that the principles underlying this organisation are similar to those we discuss here for stress-induced senescence.

## Results

### A Model for Interphase Chromosomes that Incorporates Lamina-Mediated Interactions

Most previous polymer models for interphase chromosomes have focussed on inter-and intra-chromosomal interactions and have largely neglected lamina-associated constraints on chromatin folding [6, 11, 23, 24]. In contrast, here we developed a polymer model which incorporates lamina-heterochromatin interactions to dissect the effects of lamina tethering on chromosome folding and nuclear positioning. We performed Brownian dynamics simulations which evolve the equations of motion of chromatin segments within a realistic viscous environment and subject to effective potentials modelling steric interactions and protein-mediated attraction [4, 7, 23, 24].

#### The Chromosome

LADs form large continuous blocks of chromatin which are strongly enriched in HC marks [25–28]. To account for NL-mediated interactions, we coarse-grained human chromosomes into a semi-flexible chain made of beads representing 10 kb, each set to represent either HC or EC based on the enrichment of H3K9me3 (from ChIP-seq) and LADs signal (from DamID) in the corresponding genomic region [19] (see Methods for details, and Fig. **1**A). At this resolution, a simulation of the whole genome would require about 600,000 beads, resulting in simulations too long to explore a large parameter space. In addition, the conformations we seek to explore, like SAHF, have been shown to operate on a chromosomal or smaller scale [29]. Thus, to render the simulations more computationally feasible, we considered only one chromosome, human chromosome 20, as it contains large regions of both active and inactive epigenetic marks [19], and it displays a moderate tendency to be near the nuclear periphery [30]. We considered a realistic nuclear chromatin density by performing our simulations within a box of linear size *L* = 2 *μ*m and with periodic boundaries in the *x*-*y* directions and confined in the *z* direction, so to mimic a small portion of nucleus near the NL (see Fig. **1**A). We emphasise that our framework may in principle also be employed to simulate the whole human nucleus.

**Figure 1.**
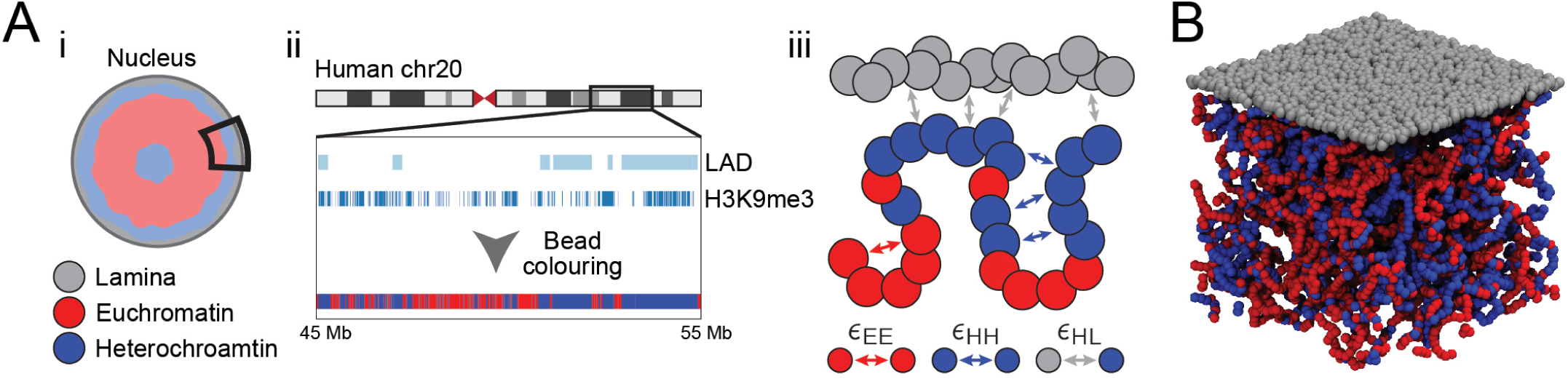
polymer model for lamina-mediated chromosome organisation in different cell states. (A) (i) A subsection of the nuclear periphery and human chromosome 20 were simulated. Chromatin was modelled as a semi-flexible bead-spring chain with red beads representing euchromatin (EC) and blue beads representing heterochromatin (HC). The NL was represented as a layer of static beads (grey). (ii) Chromatin beads were labelled as HC if the corresponding genomic region is enriched in H3K9me3 and/or LADs; all other beads were labelled as EC. (iii) EC and HC beads can interact with beads of the same kind with interaction strength *ϵ*_EE_ and *ϵ*_HH_, respectively. HC beads can also interact with the NL beads with interaction strength *ϵ*_HL_. (B) A simulation snapshot of the model when the HC-HC and HC-NL interactions are weak.

#### The Lamina

The NL was modelled by adding a thin layer (about 50 nm) of randomly positioned beads representing lamin proteins at the top of the simulation box (see Fig. **1**A,B). The interaction between HC and NL is mediated by a variety of proteins [13, 26, 31–33]. One anchor protein between NL and HC is the lamin B receptor (LBR), which has been shown to play an important role in forming the peripheral HC [32] and interacting with HP1 [31, 34]. Evidence for these protein-protein interactions and the fact that LADs are largely comprised of HC [25–28] were accounted for by setting an effective attraction between HC and NL beads via a phenomenological potential with strength given by the energy *ϵ*_HL_ (see Methods).

HP1 has been shown to dimerize *in vitro* and *in vivo* and is thought to mediate HC compaction [34–36]. In line with observations on HP1 dimerization [37] and previous modelling of intra-chromatin folding [6, 7, 38, 39], we hypothesised that HP1 mediates HC-HC interactions and thus set self-association interaction between HC beads via the same potential as for HC-NL interaction but with a strength given by the energy *ϵ*_HH_. In contrast, EC was assumed to display no tendency to self-associate, so we only included steric interactions (*ϵ*_EE_) for those beads; the result is EC displaying the typical open conformation of transcriptionally active chromatin [40, 41].

In contrast to more elaborate, many-parameter models that have been considered in the past, the behaviour of our model is governed by two key parameters, *ϵ*_HH_ and *ϵ*_HL_, which can be independently varied to explore the phase space of chromatin organisation in the presence of NL.

#### The Observables

To quantitatively characterise the organisation of the steady states displayed by the system for different combinations of the parameters (*ϵ*_HH_, *ϵ*_HL_), we computed the degree of polymer adsorption as the fraction of beads attached to the NL in steady state, *ψ*, and the overall polymer compactness as the local number density of beads, *ρ*. The former is defined as and the latter as

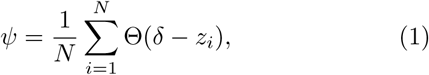

and the latter as

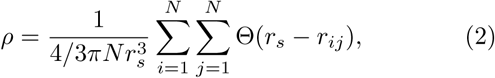

where the sums run over the *N* polymer beads in our simulation, Θ(*x*) = 1 if *x* ≥ 0 and 0 otherwise, *z_i_* is the distance of the *i*-th bead from the NL, and *δ* = 150 nm is a threshold to determine whether a bead is adsorbed to the NL. The local density *ρ* is computed within a sphere centred on bead *i* with radius *r_s_* = 250 nm and averaged across all beads in the simulation. We note that choosing different, reasonable values for *δ* and *r_s_* does not qualitatively affect the results presented below.

### Heterochromatin and Lamina Interactions are Sufficient to Capture Chromosomal Conformations Resembling Growing, Senescent and Progeroid Cells

By performing stochastic polymer simulations across the parameter space (*ϵ*_HH_, *ϵ*_HL_), we discovered that the chromatin fibre exhibits four qualitatively distinct organisations, or phases, corresponding to the four possible combinations of adsorbed/desorbed states (associated with high/low values of *ψ*) and collapsed/extended polymer conformations (associated with high/low values of *ρ*, see Fig. **2**A).

**Figure 2.**
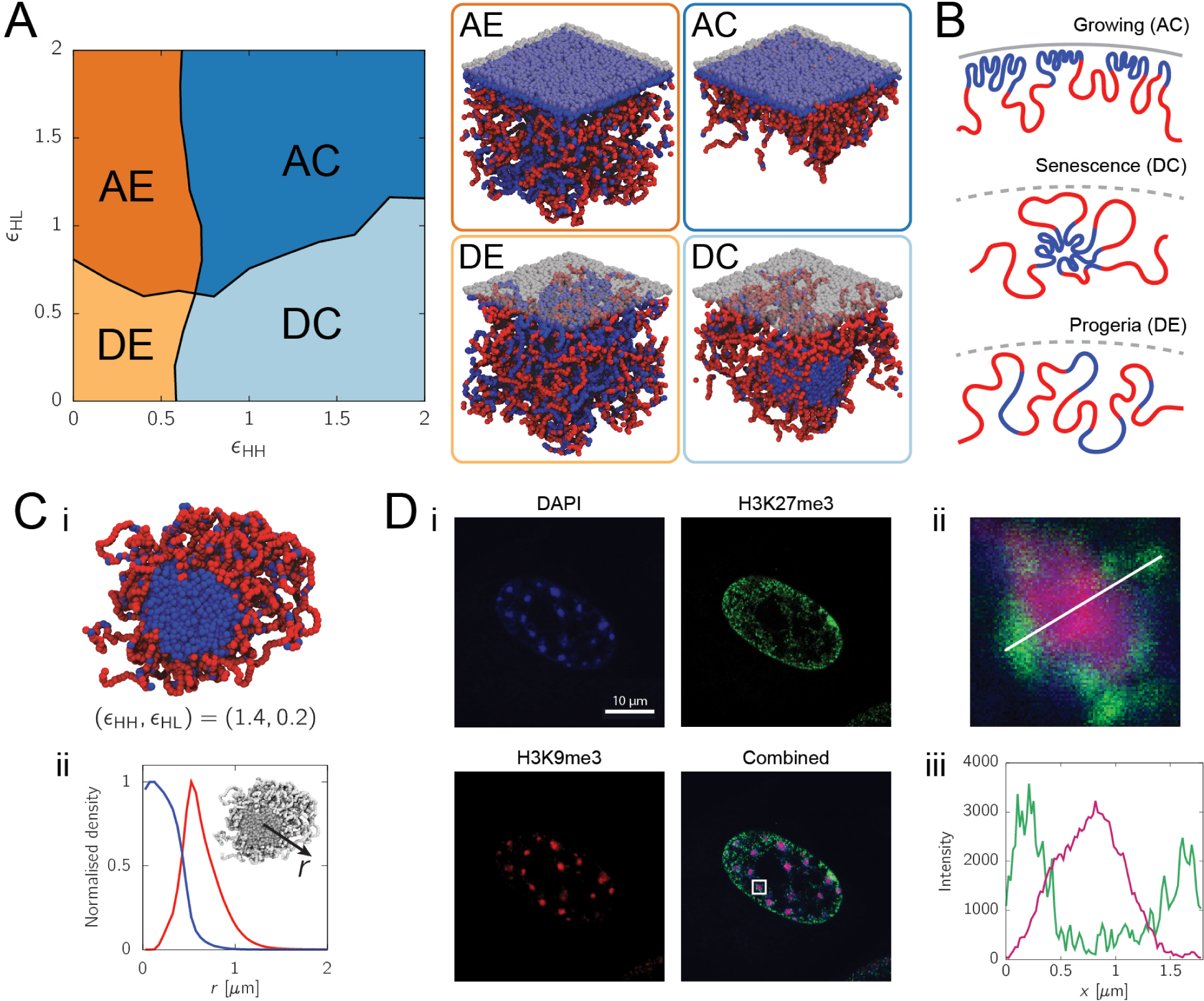
Variation of the two model parameters reproduces behaviour in growing, senescent and progeroid cells. (A) *Left* : Phases of the simulation model, computed based on two observables: the fraction of beads in contact with the lamina *ρ* (for adsorption) and the local number density of beads *ψ* (for compactness). Measuring these two parameters shows that there are four phases: adsorbed extended (AE), adsorbed collapsed (AC), desorbed extended (DE), and desorbed collapsed (DC). Phase boundaries are drawn based on the thresholds *ψ_c_* = 0.15 and *ρ_c_* = 0.2 (see SI for more details). *Right*: Simulation snapshots of the four phases. (B) Cartoons of chromatin structures for cells in growing, senescent and progeroid conditions, respectively. (C) (i) Cross-section view of a simulation snapshot corresponding to the DC/senescent phase. (ii) Corresponding density profiles of HC and EC as a function of distance from the centre of the globule, *r*. (D) (i) Confocal images of chromosomes in senescent cells with H3K9me3, H3K27me3 and DAPI staining (see Methods). (ii) Zoomed-in view of a SAHF corresponding to the white square in the combined image in (Di). (iii) Corresponding intensity profiles for H3K9me3 and H3K27me3 along the white line in (Dii).

Three of the four phases display morphologies remarkably similar to those of healthy (growing), senescent or progeroid mammalian cells (see cartoons in Fig. **2**B). Specifically, the two *ψ* > 0 phases which display a layer of HC adsorbed to the NL (the adsorbed collapsed, or AC, phase with *ρ* > 0 and the adsorbed extended, or AE, phase with *ρ* ≃ 0) are qualitatively similar to healthy, growing cells. However, the AE phase displays a significant amount of HC intermixed with EC, which is not observed in conventional mammalian cells [42, 43]; thus, we identified the AC phase as the closest representative of a cell in the growing state. The two *ψ* = 0 phases where no chromatin is adsorbed to the NL (the desorbed collapsed, or DC, phase with *ρ* > 0 and the desorbed extended, or DE, phase with *ρ* ≃ 0) display features akin to those observed in senescent and progeroid cells. Particularly, the DC phase chromosome structure markedly resembles the nucleus of a senescent cell in which HC self-associates into large, SAHF-like bodies surrounded by a corona of EC [20, 43] (Fig. **2**C,D). The DE phase instead displays phenotypes reported in progeroid cell nuclei, including the loss of peripheral HC [44–46] and a large degree of mixing of chromatin regions with active and inactive epigenetic marks [47].

Within our simulations, the DC/senescent phase is associated with extensive HC-HC interactions and weak HC-NL interactions. Nevertheless, there have been conflicting views on the role played by HC-HC interactions in the process of SAHF formation. On one hand, it has been suggested that H3K9me3 levels, which present the binding site for HP1 proteins on the chromatin, remain unchanged in senescence. On the other hand, a qualitative upregulation of one HP1 protein (HP1 beta) has been described [18]. To clarify this, we quantitatively assessed the changes in proteins able to mediate HC-HC interactions. We performed a chromatin fractionation for growing and senescent cells followed by mass spectrometry (see Fig. **3** and Methods). Indeed, we can report a consistent upregulation of HP1 proteins and macroH2A, which has been previously implicated in SAHF formation and heterochromatin compaction [48]. Therefore, as suggested by the simulations, SAHF formation might be driven through HC-HC interactions. Additionally, our mass spectrometry data show that high mobility group box (HMGB) proteins are downregulated in senescent cells, similarly to previous reports showing that HMGB2 is lost when cells enter senescence [49].

**Figure 3.**
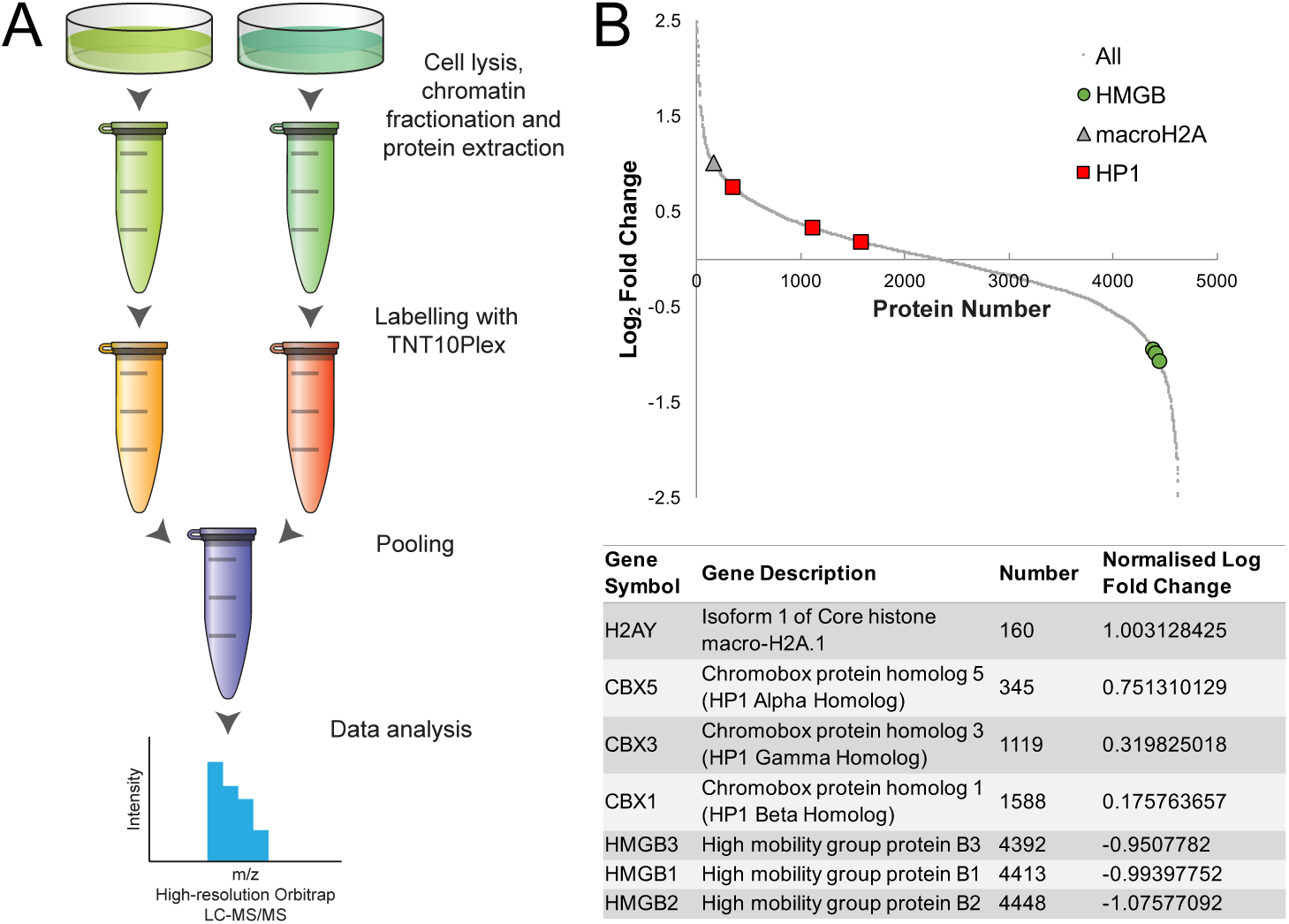
Chromatin fractionation and mass spectroscopy experiments investigating the change in protein abundance between growing and senescent cells. (A) A schematic representation for chromatin fractionation from different samples followed by identification and quantitation of proteins based on high-resolution Orbitrap LC-MS/MS. (B) *Top*: A graph showing the normalised ratio of protein abundance between growing and senescent cells. Proteins are numbered from the largest to the lowest ratio. *Bottom*: A table listing the normalised ratios for macro-H2A, HP1 and high mobility group proteins.

Given the simplicity and low-dimensionality of the parameter space scanned by our simulations, it is remarkable that our model can capture key qualitative features of chromosomal organisation in a range of different cell states. In light of this, we concluded that the two key ingredients of our model, HC-HC and HC-NL interactions, must be the major driving forces of chromatin folding in these cell states and can guide the dynamical re-organisation of the nuclear architecture upon transition to different physiological and pathological conditions.

### Phase States Resembling Senescence and Progeria Predict Global Differences in the Network of Chromatin Contacts

To address whether our polymer model could quantitatively capture the changes in the pattern of chromatin contacts in senescent and progeroid cells measured via HiC experiments [20], we performed further simulations in which we included a weak self-attraction between EC segments (*ϵ*_EE_ = 0.4*k_B_ T*) to more closely mimic the formation of A compartments [2]. We also explicitly modelled the centromere, covering a region between 26.4-29.4 Mb along chromosome 20, as a separate, self-associating type of bead (also with an interaction energy *ϵ*_CC_ = 0.4*k_B_ T*). From these simulations, we computed the network of contacts between beads in different phases of the system. Circos plots of this interaction network (Figs. **4**Ai,Ci) readily show that while senescent simulations display more longrange contacts with respect to growing cells, those for progeria show a lack of distal contacts. The same qualitative behaviour is observed in similar plots for growing, senescent and progeroid cells based on HiC data (obtained from Refs. [20] and [47] respectively; Figs. **4**Bi,Di).

**Figure 4.**
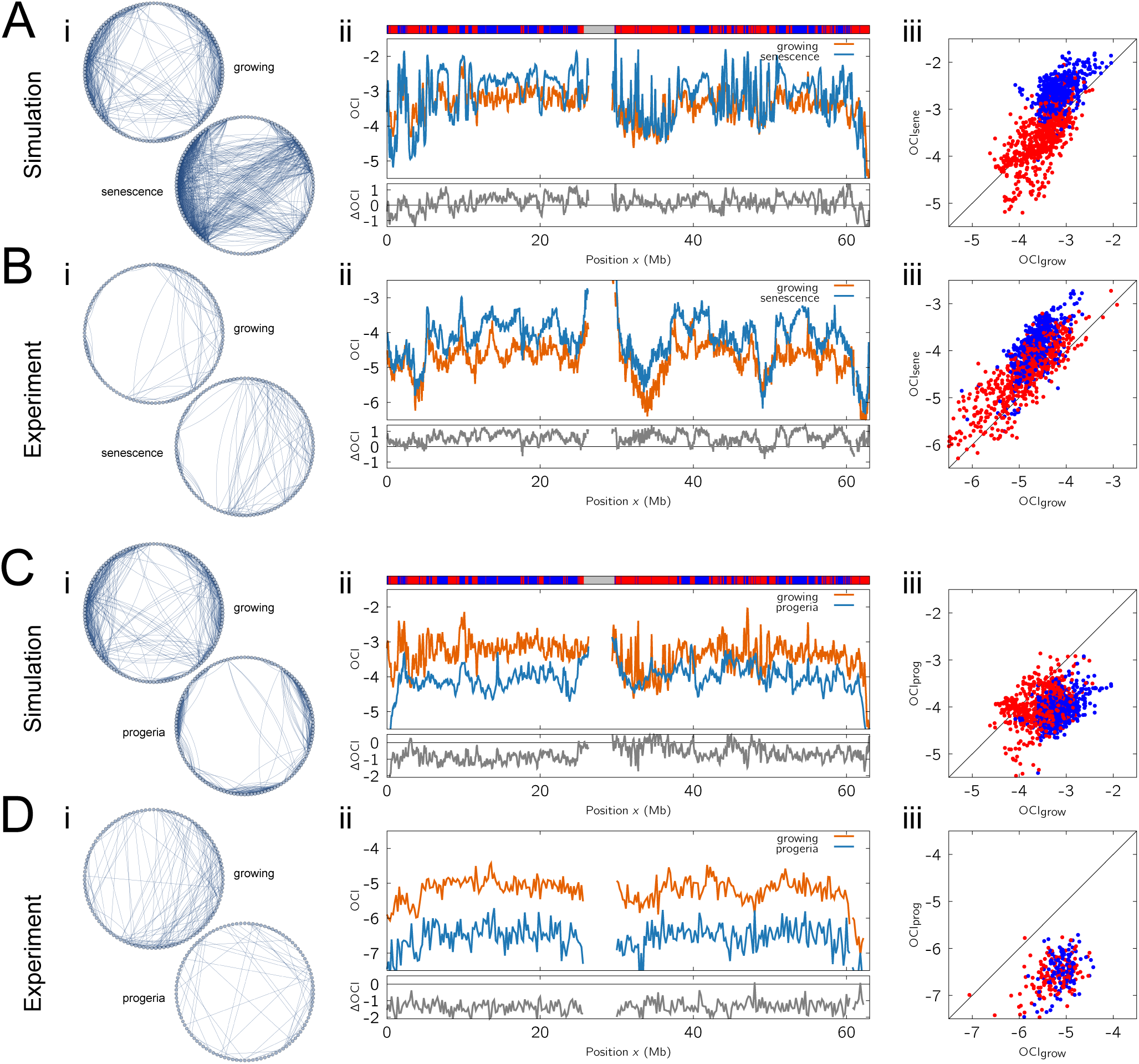
Chromatin contact network in (AC) growing, (DC) senescent and (DE) progeroid state. For simulations, we used (*ϵ*_HH_, *ϵ*_HL_) = (1.4, 1.8) for growing, (1.4, 0.2) for senescence, and (0.4, 0.2) for progeria. In (A) and (B), we compare the contact patterns of growing and senescence phase in our model to those in HiC data from Ref. [20]. In (C) and (D), we compare our results for growing and progeria to those in HiC data from Ref. [47]. The left panel (i) shows arc diagrams which give visual representations of the contact network within the chromosome in different phases. The nodes represent a subset (10%) of the chromatin bins in the order in which they appear along the chromosome. Connections drawn between bins are those with a contact frequency that is within the top ∼10% of all contacts. The middle panel (ii) reports the OCI values for the indicated phases and their difference (ΔOCI). The track at the top indicates the chromatin state of each bin (red for EC, blue for HC and grey for centromere). The right panel (iii) shows scatter plots of the OCI values of each bin in the two phases considered in the other panels. The colour of each point indicates the chromatin state of the corresponding bin.

To quantitatively characterise the change in the network of contacts, we generated simulated HiC maps by measuring the contact probability of two beads over 20 equilibrated polymer conformations [7]; we then calculated the ratio of distal to local contacts at each chromosome region, also known as the “open chromatin index” (OCI) [20] (with threshold set at 2 Mb, see Methods). The difference in OCI, ΔOCI, readily quantifies the changes in the network structure upon transitioning from the growing to the senescent state [20].

From our simulation results, we could detect a dramatic change in the network of chromatin contacts in the senescence (DC) and progeria (DE) states compared to growing (AC) state. More specifically, the transition from growing (AC) to senescent (DC) phase is characterised by an overall positive and statistically significant ΔOCI (two-sample Kolmogorov-Smirnov (KS) test: *D* = 0.31, *p* < 10^−4^; a larger *D* means that the two samples are drawn from more separated distributions; see Fig. **4**Aii). This finding implies that, as qualitatively shown in the Circos plots, there is a substantial increase in distal contacts. In contrast, our simulations predicted an overall negative ΔOCI associated with the growing-progeria transition, i.e. comparing the AC phase to the DE phase, which entails a chromosome-wide loss of distal contacts (two-sample KS test: *D* = 0.65, *p* < 10^−4^; Fig. **4**Cii).

We next asked whether the predictions from our simulations could be verified using the HiC data. To this end, we computed the OCI for chromosome 20 (excluding inter-chromosomal interactions). The signal shows the same quantitative behaviour as that from our simulations, i.e. senescent cells gain distal contacts (Fig. **4**Bii) whereas progeroid cells lose them (Fig. **4**Dii). In both cases the difference is statistically significant: for senescence we measure *D* = 0.47, *p* < 10^−4^, and for progeria *D* = 0.88, *p* < 10^−4^.

It is remarkable that while both senescence and progeria cell states display a global loss of chromatin-lamina interactions, they show an opposite change in chromatin contacts captured by the OCI. We reasoned that this difference may be associated with the appearance of segregated HC compartments in senescence but not in progeria. To verify this hypothesis, we constructed scatter plots showing the OCI value of each bead, colour-coded based on its epigenetic state (Figs. **4**Aiii-Diii). These plots reveal that the change in OCI associated with the growing-senescence transition is stronger in HC-rich regions compared with EC-rich regions (blue points appear further from the diagonal; Figs. **4**Aiii,Biii). This trend was not found when comparing growing with progeroid cells (Figs. **4**Ciii,Diii). This result is consistent with the finding that GC-poor isochores exhibit a larger change in their interaction network between growing and senescence [20], and it further illustrates the fundamental role played by the competition between HC-HC and HC-NL interactions in shaping the nuclear landscape and its re-organisation upon changes in physiological conditions. Furthermore, this finding constitutes compelling evidence that SAHF, present in senescent cells but notably absent in progeria, may be mediating the emergence of long-range chromatin contacts by forming a polymer-polymer phase separation between HP1 and HC-rich chromatin [7, 10, 50, 51].

To further quantify the agreement between simulation and experiment, we calculated the Pearson correlation coefficient for ΔOCI as a function of position along the chromosome. For the growing versus senescence case, we found a correlation of *r* = 0.37 (*p* < 10^−4^) between simulation and experiment. For the growing versus progeria case, the correlation was *r* = 0.21 (*p* < 10^−3^).

### Observed Changes in Chromatin Network Corresponds to a Change in the Polymer Metric Exponent

To rationalise the observed opposite change in distal contacts in senescent and progeroid cells, we considered the contact probability of two chromatin segments as a function of their genomic distance *s*, *P*(*s*), which can be extracted from HiC maps [52]. Classic results from polymer physics predict that in three dimensions the contact probability can be described by the scaling *P*(*s*) ∼ *s*^−3*v*^ [21, 53], where *v* is known as the metric exponent of a polymer. For large enough chains, *v* also determines the scaling of the typical polymer size with its length i.e., *R* ∼ *N^v^*, where *N* is the number of segments [40].

Values of *v* have been determined for different types of polymer conformations and solvent conditions [54, 55]. By analysing the conformations assumed by the simulated chromosome in the different phases, we may associate the senescent (DC) phase with crumpled globules, i.e. *v*DC = 1/3 [55], whereas the progeria (DE) phase is consistent with a melt of self-avoiding polymers, i.e. random walks (RW) with *v*DE = 1/2 [54]. At the same time, since the growing (AC) phase is collapsed but also adsorbed at a surface (the NL), we reasoned that its associated metric exponent must be strictly larger than that for a crumpled globule but smaller than that for a RW, i.e. *v*DC < *v*AC < *v*DE. This mapping implies that the growing-senescence and growing-progeroid transitions are associated with a change in the metric exponent *v*, which in turn leads to a change in the contact probability *P*(*s*). Then, because contact probabilities are normalised, a smaller value of *v* (as is the case, for instance, for the DC/senescent phase with respect to the growing phase) leads to a shallower decay in contact probability, hence to a shift favouring non-local over local contacts.

More quantitatively, we can thus compute the OCI as

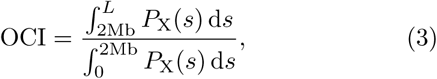

where X can either be AC, DC or DE, and *L* is the total length of the chromosome. Given that *P*_AC_(*s*) ∼ *s*^−3*v*_AC_^ decays more quickly than *P*_DC_(*s*) ∼ *s*^−1^ and more slowly than *P*_DE_(*s*) ∼ *s*^−3/2^, Eq. (3) predicts that the change in OCI must be positive upon triggering a senescent state and negative upon entering a progeroid state, as found in both simulation and experiment.

### LADs Display Cell-to-Cell Variability

Lamin association is a major regulator of nuclear architecture, and LADs cover more than 30% of the humangenome [13]. Yet, each cell has only a limited amount of surface that is available for chromatin to interact with. For this reason it has been conjectured, and then shown, that LADs display cell-to-cell variability, appear stochastically and are not conserved in daughter cells [27, 28]. By measuring the adsorption of beads onto the NL in single, independent polymer simulations, which we associated with individual cells, we found that this stochasticity is intrinsically captured within our model (Fig. **5**A). Remarkably, even though all HC beads are in principle able to be adsorbed at the NL, we found that only about 50% of them are adsorbed in any one simulation. Interestingly, this corresponds to a fraction just below 30% of the whole chromosome, similar to the overall LAD coverage observed experimentally (although it is possible that this match is due to a fortunate parameter choice for the modelling of the NL).

**Figure 5.**
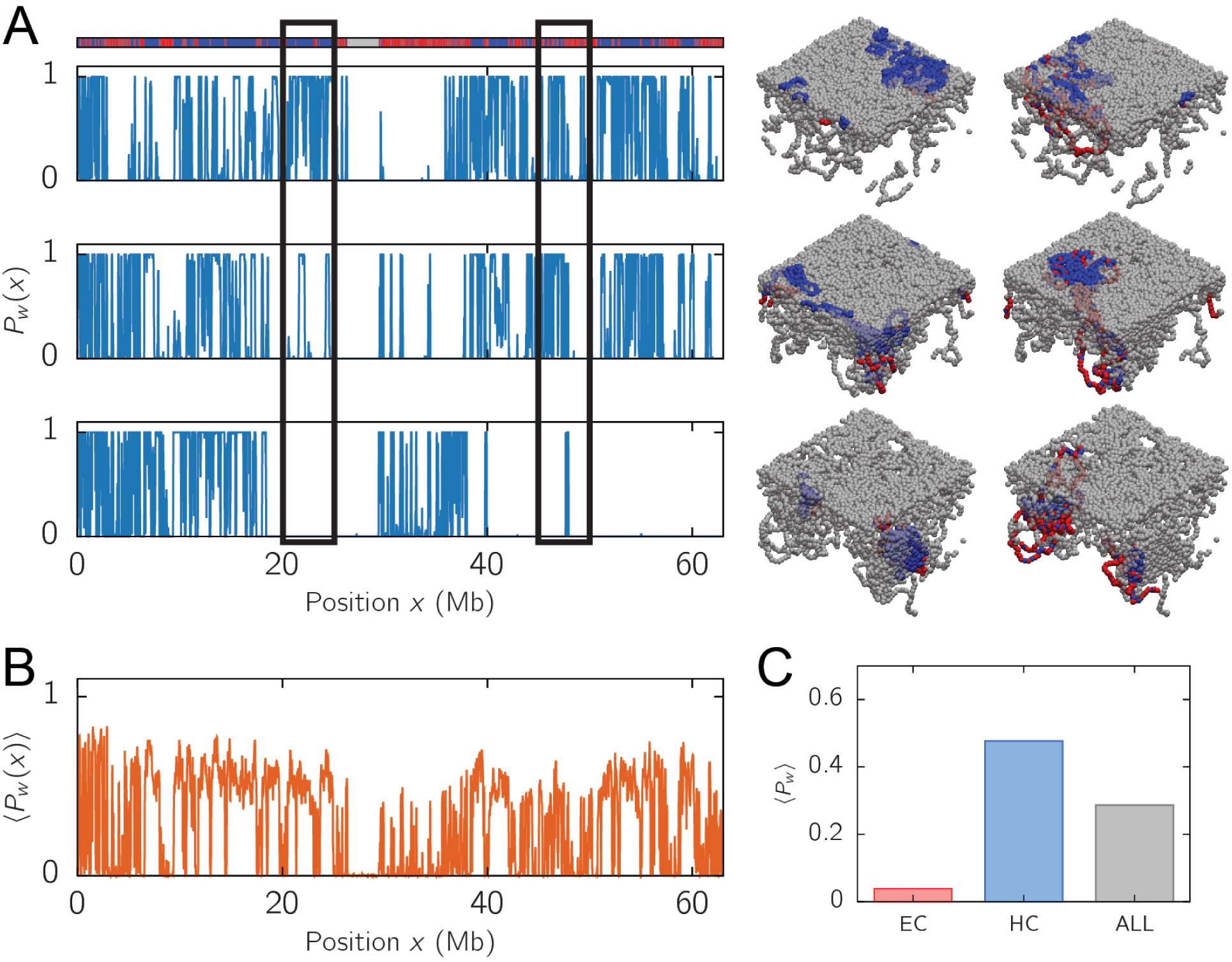
Probability of contacting the NL. (A) *Left*: Plots showing the contact probability of each bead with the lamina *P_w_*(*x*) = (Θ(*δ* – *z*(*x*)))*t* for three simulation runs in the AC (growing) phase (*ϵ*_HH_ = 1.4, *ϵ*_HL_ = 1.8). The *x*-axis shows the genomic position corresponding to each bead. The track at the top indicates the chromatin state along the polymer (red for EC, blue for HC and grey for centromeric region). *Right*: Snapshots of the simulation runs colouring only the beads within the two highlighted regions on the left panel (rectangular boxes). These figures reveal that the same chromatin segment can reside in different positions relative to the NL in different runs, indicating that LADs associate with the NL stochastically. (B) The contact probability of each bead with the NL averaged over 20 simulation runs. (C) The average contact probability with the NL for EC, HC and all beads.

Once averaged over all simulation replicas (equivalent to averaging over a population), we discovered that the adsorption probability for each bead differs noticeably from those in single simulation runs (Fig. **5**B). This result emphasises that population averaged information on chromatin conformation, such as HiC, may not fully reflect the conformation assumed in single cells [56].

It is also intriguing to note that there is a non-zero probability for a EC bead to be adsorbed to the NL, even though there is no explicit attractive interaction between the two in the model. We reasoned that this observation originates from the chromatin context in which a given EC bead is embedded – i.e. a EC-rich segment within a large HC-rich chromosome region is likely to be “dragged along” and co-adsorbed onto the NL (see also simulation snapshots in Fig. **5**A). This result indicates the importance of considering the context of neighbouring chromatin when determining the spatial location and function of a specific locus.

### A First-Order-Like Phase Transition Separates Growing and Senescent States

After showing that our polymer model can capture many complex features observed in growing, senescent and progeroid cells, we employed it to obtain further insight into the nature of the transition between different states of a cell. This is of fundamental interest to polymer physics and has biological relevance, as the nature of the transition determines the stability of the growing and senescent states upon external perturbations, such as an abrupt change in the number of active HP1 or lamin proteins.

To characterise the nature (order) of the phase transition between the growing to the senescent state, we asked whether this transition is associated with hysteresis [51]. To this end, we recorded the instantaneous value attained by the adsorption parameter *ψ* as we slowly varied the strength of HC-NL interaction, *ϵ*_HL_, between two values known to be deep into the respective phases (see Fig. **6**A). By starting with a large value for *ϵ*_HH_ (AC phase), slowly reducing it to a low value (DC phase), before reversing the process and slowly increasing it again, we discovered that the change in *ψ* does not follow the same pattern in each direction. That is to say, the phase transition does not occur at the same critical value of HC-NL interaction strength, and the system is thus history-dependent (Fig. **2**).

**Figure 6.**
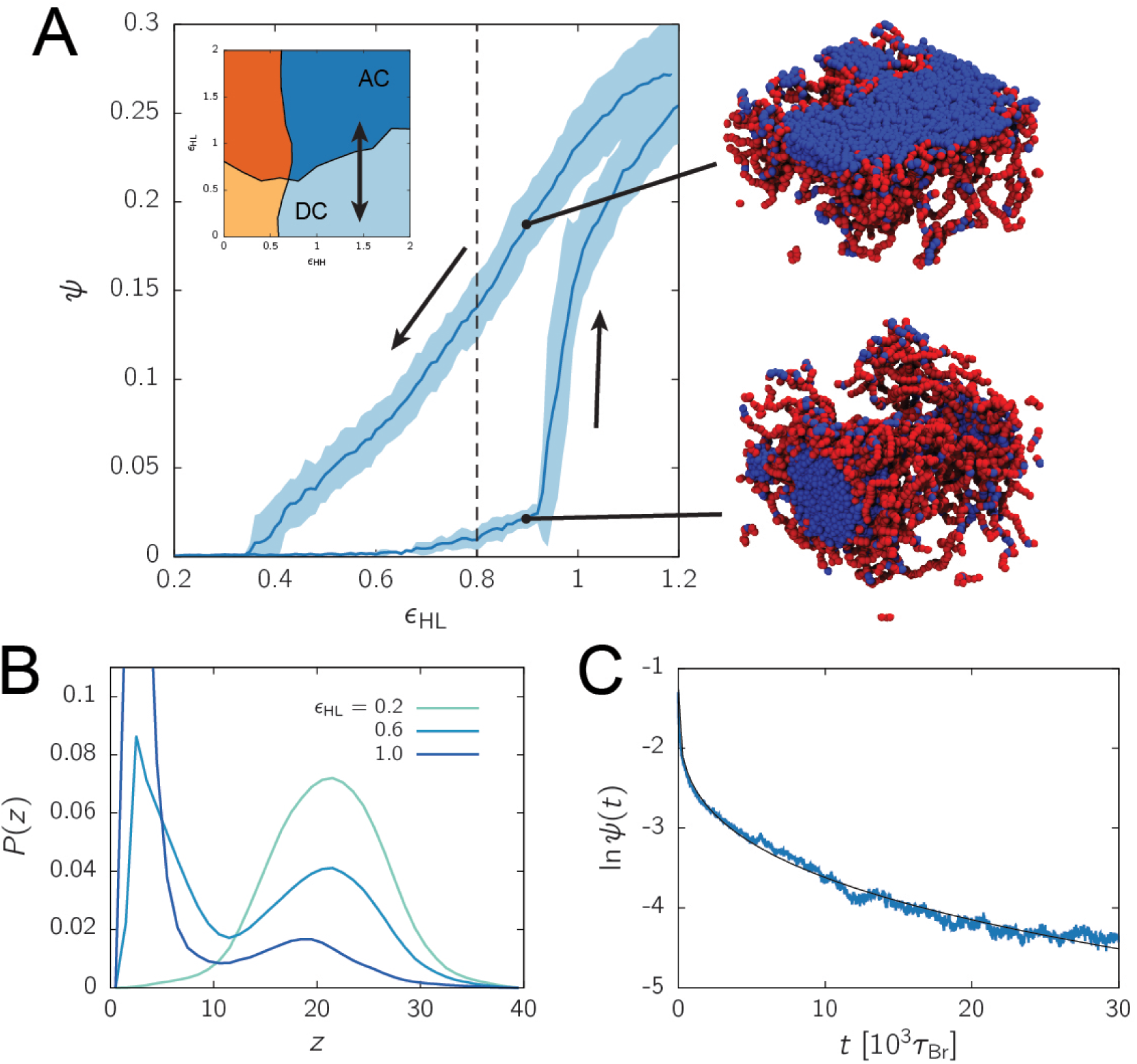
A first-order-like transition between the AC (growing) and DC (senenscent) phase. (A) The fraction of beads in contact with the NL, *ψ*, averaged over five simulation runs in which we fix (*ϵ*_HH_ = 1.4 and vary *ϵ*_HL_ between 0.2 and 1.2 with 10^6^ Brownian time steps in each direction (the path in the phase diagram is shown in the inset). The shaded area around the curve reports the standard deviation of the mean. Hysteresis occurs in the region *ϵ*_HL_ ∼ 0.4-0.9. The snapshots on the right show that the system can be in either the AC (growing) and DC (senescent) phase at *ϵ*_HL_ = 0.9 depending on its history. (B) The probability of finding a HC bead at distance *z* from the wall for *ϵ*_HL_ between 0.0 and 1.0 with *ϵ*_HH_ = 1.0 at equilibrium. A bimodal behaviour is found when *ϵ*_HL_ ∼ 0.6. (C) The change in *ψ* with time *t* in a log-linear plot after reducing *ϵ*_HL_ from 1.2 to 0.4 instantaneously. The black line shows a stretch-exponential fit *f*(*t*) ∼ exp(–*αt^β^*) with *α* = 0.16 and *β* = 0.29.

Such a hysteresis cycle is a well-known hallmark of a first-order phase transition between two equilibrium phases and has a number of biologically relevant consequences. For instance, the passage from growing to senescence is believed to be highly irreversible. This permanent growth arrest is a mechanism by which cells can prevent stresses, e.g. induced by an oncogene or by DNA damage, from inflicting further harm to the organism. It is therefore important for senescent cells to remain stable and not to return to the proliferating state. A first-order transition, with strong hysteresis, is fully consistent with this irreversibility. It suggests that structures found in senescence, in particular SAHF, are stable under perturbations and that there are physical mechanisms inhibiting senescent cells from inappropriately turning into growing cells. Further, a first-order transition means that the change from one state to the other is abrupt and occurs in a step-wise manner within a short window of parameter space. This can be observed by measuring the distribution of HC beads at distance *z* from the NL, *P*(*z*) (Fig. **6**B). As the HC-NL interaction strength is decreased, the system jumps from an adsorbed state to a desorbed state populated by SAHF-like structures.

We reasoned that such an abrupt transition can be explained by arguing that the EC-rich chromosomal regions surrounding the SAHF in the senescent (DC) state (Fig. **2**) provide a large entropic barrier that needs to be overcome in order for the HC beads to be adsorbed at the NL [57]. In other words, even if the HC beads within the SAHF can bind the NL, a large scale rearrangement of the surrounding halo of EC beads is required for the HC and NL beads to come into contact. This stabilises the senescent DC phase. A similar argument suggests that the growing (AC) phase is also stable to perturbations as the transition into the DC phase entails a large-scale rearrangement to form the SAHF surrounded by the EC corona, and this rearrangement is likely associated with another free energy barrier.

LADs detachment from the nuclear periphery is a common phenotype in cellular senescence; however, the dynamics of such process is not well understood. Here, we employed our model to provide a prediction on how LADs separate from the lamina as cells enter senescence (for instance, as a response to DNA damage). Specifically, we monitored the fraction of chromatin segments close to the NL over time after the HC-NL interaction, *ϵ*_HL_, was reduced abruptly, mimicking the loss of lamina interactions at the onset of senescence (Fig. **6**C). We observed that the decay is non-exponential. Previous polymer studies reporting this phenomenon demonstrated that such a decay trend can arise if polymer segments are desorbed in a co-operative manner [58], or if the desorption process is limited by the diffusion of chain segments away from the surface [59].

## Discussion and Conclusions

In this work, we proposed and studied a polymer model for nuclear organisation which explicitly takes into account chromatin-lamina interactions. A key result is that by modifying only two parameters – the HC-HC self-interaction and the HC-lamina (HC-NL) interaction – our model displays four possible distinct phases, three of which recapitulate nuclear architectures which are biologically relevant. The adsorbed collapsed (AC) phase corresponds to the morphology of conventional growing cells, where HC forms a layer close to the lamina, and HC-HC interactions drive (micro)phase separation of HC and EC compartments. In stress-induced cellular senescence and progeria, chromatin-lamina interactions are disrupted; hence their corresponding nuclear structure is captured by desorbed phases, where HC-NL interactions are weak. More specifically, the phenotype of cellular senescence, which is associated with extensive HC foci, corresponds to our desorbed collapsed (DC) phase, where HC-mediated interactions still drive clustering (phase separation) but HC is not bound to the lamina, whereas the desorbed extended (DE) phase is consistent with the sparse chromosome organisation of progeroid cells.

This classification is in line with qualitative expectations from existing biological models, but the link we make here to polymer physics allows us to further quantitatively explain additional features of nuclear organisation in growing and senescent cells which were previously mysterious. Viewing the stress-induced entry to senescence as a phase transition (between the AC and DC phases) naturally leads to the prediction that intra-chromosomal contacts should become less local and longer-range. Theories of polymer looping predict that the probability of contact decays more steeply with distance along the backbone (in genomic distance) in more swollen, or open, polymer configurations than in more compact, or globular, conformations. This trend matches existing experimental measurements of the “open chromatin index”, via HiC, which find a different network of chromatin contacts in growing and senescence cells, with more distal contacts typically appearing in the latter. The same argument suggests that, in contrast, when a growing cell becomes progeroid, contacts should become more local, as the DE phase is less compact than the AC phase. Notably, by re-analysing HiC data for HGPS cells [47], we found that similar trends occur *in vivo*.

The finding that the inter-chromatin contact network changes in opposing ways in progeroid and senescent cells (favouring more local contacts in the former, and more distal in the latter) might seem at first surprising, as progeria is normally viewed as the first step towards senescence [20]. However, our model shows that this result is a consequence of the fact that progeroid, growing and senescent architecture correspond to different thermodynamic phases for the melt of interphase chromosomes in the nucleus.

Our polymer physics framework also yields additional predictions: some of these conform well with existing experimental evidence, while others could be tested by new experiments in the future. For example, our model clearly shows that lamina adsorption is stochastic: different HC domains bind the lamina in different simulations, and not all HC is absorbed. This is in line with single-cell DamID experiments which show that not all LADs contact the lamina in every cell, and that the selection of which LADs are bound to the nuclear periphery in any given cell is largely random [13]. Also, our model predicts that the transition between growing and senescent cells should be first-order-like, so that each state, once established, should be metastable even in parameter space regions where it is normally unstable. This provides an appealing mechanism to explain the remarkable stability of senescent cells: once SAHF form, the cell virtually reaches a thermodynamic dead end (i.e., it is unlikely to ever re-enter the normal cell cycle of proliferating cells, to grow and divide again). Our results suggest that the enhanced stability of the senescent phenotype is largely due to the active EC layer which provides a barrier preventing the HC core of each of the SAHF from reaching the lamina.

As for predictions which could be tested in the future, we suggest that these would entail studying the dynamics of chromosome desorption following the entry into senesence. First, we expect that SAHF should form by coarsening, and the associated growth laws could be compared between simulations and live cell microscopy. Second, we suggest that monitoring the amount of adsorbed chromatin as a function of time will be of interest: this will test our predictions that its decay is non-exponential, and may uncover spatiotemporal correlations between LAD dynamics. It would also be instructive to assess the role of the confining geometry in determining the final morphology of the system. In stress-induced senescence, desorption leads to the formation of a SAHF for each chromosome. In the rod cells of nocturnal mammals, HC also detaches from the lamina but forms a single, larger aggregate [60]. It is attractive to think that this may be due to the smaller size of retinal cells with respect to senescent cells: does the enhanced confinement push HC into larger clusters? It might be possible to perturb the geometry of senescent cells to further address the role of nuclear geometry in chromosome architecture: this question could be asked both experimentally and in simulations. Finally, our work shows that the phase separation of HC and EC couples to nuclear topography and architecture in a profound way, and it would be important to find out whether the phase-separated and layered organisation in growing cells offers any functional advantage with respect to the organisation in other cell states.

## Materials and Methods

### Fluorescence Imaging of Senescent Cells

We cultured IMR90-ER:RAS cells, which expressed a chimeric fusion protein upon induction by 4-hydroxytamoxifen (4OHT), in 100 nM 4OHT in DMEM 10% FCS at 37°C atmospheric oxygen. Growth arrest was triggered after 7 days of 4OHT treatment and cells became senescent. Cells were plated on gelatin-treated coverslips and allowed to attach to the surface overnight. Cells were fixed in 4% paraformaldehyde (in 1×PBS) for 15 min and washed three times. Fixed cells were permeabilised using 0.2% Triton X/PBS for 5 min at RT. Primary antibodies (Anti-Histone H3 (tri methyl K9) [Ab8898, Abcam] plus Anti-Histone H3 (tri methyl K27) [Ab6002, Abcam], or (Anti-Histone H3 (tri methyl K36) [Ab9050, Abcam] plus Anti-Histone H3 (tri methyl K9) [05-1242-S, Millipore]) were diluted in 1:1000 ratio (1×PBS) and incubated on the cells for 45 min at RT. The cells were washed with the blocking solution PBS-T (0.1% Tween in 1×PBS) for 30 min, followed by the incubation with fluorophore conjugated secondary antibodies (Alexa Fluor^®^ 488 anti-rabbit and Alexa Fluor^®^ 555 anti-mouse) for 45 min in darkness at RT. After secondary incubation, the cells underwent additional washes with PBS-T for another 30 min before being dried and mounted with Vectashield antifade mounting medium containing 4’-6-diamidine-2-phenyl indole (DAPI). We used the Nikon’s A1R point scanning confocal laser-scanning microscope for capturing immunofluorescent images. To optimise the quality of image acquisition, the Nikon’s CFI Plan Fluor 60× oil objective lens was selected to acquire images. The triple band excitation DAPI-FITC-TRITC filter was used to detect fluorescent signals from DAPI, Alexa Fluor^®^ 488 and Alexa Fluor^®^ 555, respectively. The laser power was set at 4.00 and the detector sensitivity fixed at 80 for every fluorescence channel. The pinhole was set at 1.2 AU and the images acquired were 2048×2048 in pixel.

### Chromatin Fractionation and Mass Spectrometry

#### Chromatin Isolation

Cells were trypsinised and resuspended in 10-15 mL DMEM. Cells were counted, the same number of cells collected from each sample by centrifugation, and resuspended in cold PBS. Supernatant was removed and cells resuspended in no-salt buffer A (DTT, PMSF, Digitonin) and incubated on ice for 10 min. Cells were centrifuged and supernatant (cytosolic components) kept for further preparation. Nuclei pellet was resuspended in no-salt buffer A (DTT, PMSF) and centrifuged. Supernatant was removed and nuclei pellet resuspended in no-salt buffer B (DTT, PMSF) and vortexed occasionally on ice before centrifuging. The nucleoplasmic supernatant fraction was kept for further analysis and the remaining chromatin pellet resuspended in no-salt buffer B followed by centrifugation. Supernatant was removed and chromatin pellet resuspended in 1× Sample buffer. Suspension was vortexed and heated in boiling water for 5 min to release nuclear-bound proteins.

#### Chromatin Fractionation

Cells were harvested and centrifuged. Cell pellets were resuspended in cold lysis buffer and mixed thoroughly. Cell lysate was centrifuged and supernatant transferred to a new tube. Protein concentration was determined (BCA Protein Assay Kit). Proteins extracted were reduced, alkylated, digested overnight and labelled as described in the TMT10plex Mass Tag Labeling Kits and Reagents. Labelled samples were analysed by high-resolution Orbitrap LC-MS/MS (see the next section). Labelled peptides were identified and reporter ion relative abundance quantified.

#### TMT Labelling and Mass Spectrometry

TMT labelling was performed according to the manufacturer’s protocol (https://www.thermofisher.com/order/catalog/product/90110). 8 samples were digested and the resulting peptides were labelled with the tags 126, 127C, 128C, 129N, 129C, 130N, 130C and 131, before being combined and lyophilised.

The following LC conditions were used for the fractionation of the TMT samples: desalted peptides were resuspended in 0.1 mL 20 mM ammonium formate (pH 10.0) + 4% (v/v) acetonitrile. Peptides were loaded onto an Acquity bridged ethyl hybrid C18 UPLC column (Waters; 2.1 mm i.d. × 150 mm, 1.7 m particle size), and profiled with a linear gradient of 5-60% acetonitrile + 20 mM ammonium formate (pH10.0) over 60 min, at a flow-rate of 0.25 mL/min. Chromatographic performance was monitored by sampling eluate with a diode array detector (Acquity UPLC, Waters) scanning between wavelengths of 200 and 400 nm. Samples were collected in one-minute increments and reduced to dryness by vacuum centrifugation, before being pooled into pairs (in total, 13 paired fractions were generated). Fractions were resuspended in 30 mL of 0.1% formic acid and pipetted into sample vials.

All LC-MS/MS experiments were performed using a Dionex Ultimate 3000 RSLC nanoUPLC (Thermo Fisher Scientific Inc, Waltham, MA, USA) system and a QExactive Orbitrap mass spectrometer (Thermo Fisher Scientific Inc, Waltham, MA, USA). Separation of peptides was performed by reverse-phase chromatography at a flow rate of 300 nL/min and a Thermo Scientific reverse-phase nano Easy-spray column (Thermo Scientific PepMap C18, 2 mm particle size, 100 A pore size, 75 mm i.d. × 50 cm length). Peptides were loaded onto a pre-column (Thermo Scientific PepMap 100 C18, 5 mm particle size, 100 A pore size, 300 mm i.d. × 5 mm length) from the Ultimate 3000 autosampler with 0.1% formic acid for 3 min at a flow rate of 10 mL/min. After this period, the column valve was switched to allow elution of peptides from the pre-column onto the analytical column. Solvent A was water + 0.1% formic acid and solvent B was 80% acetonitrile, 20% water + 0.1% formic acid. The linear gradient employed was 4-40% B in 100 min (the total run time including column washing and re-equilibration was 120 min).

The LC eluant was sprayed into the mass spectrometer by means of an Easy-spray source (Thermo Fisher Scientific Inc.). All *m/z* values of eluting ions were measured in an Orbitrap mass analyser, set at a resolution of 70000. Data dependent scans (Top 20) were employed to automatically isolate and generate fragment ions by higher energy collisional dissociation (HCD) in the quadrupole mass analyser and measurement of the resulting fragment ions was performed in the Orbitrap analyser, set at a resolution of 35000. Peptide ions with charge states of between 2+ and 5+ were selected for fragmentation.

Proteome Discoverer v1.4 (Thermo Fisher Scientific) and Mascot (Matrix Science) v2.2 were used to process raw data files. Data was aligned with the UniProt human database, in addition to using the common repository of adventitious proteins (cRAP) v1.0. Protein identification allowed an MS tolerance of °20 ppm and an MS/MS tolerance of °0.1 Da along with up to 2 missed tryptic cleavages. Quantification was achieved by calculating the sum of centroided reporter ions within a °2 millimass unit (mmu) window around the expected *m/z* for each of the 8 TMT reporter ions.

### Simulation Details

#### Computational Modelling of Chromatin and Lamina

We modelled the chromosome as a semi-flexible bead-spring chain of *N* beads. Each bead represents a 10 kb chromatin segment, which is equivalent to about 50 nucleosomes and has a diameter of *σ* ∼ 50 nm. As mentioned in the main text, we chose to model human chromosome 20 (*N* = 6303). Each bead was assigned one of two possible types *q*: one for euchromatin (EC) (*q* = 1, coloured red in the figure) and the other for heterochromatin (HC) (*q* = 2, blue) (Fig. **1**Ai). Beads were coloured using ChIP-seq data for H3K9me3 and Dam-ID data for LADs available on ENCODE [19] and aligned to the GRCh37/hg19 assembly. The LAD data were from the Tig3 cell line [26]. For the H3K9me3 data, we used those from the GM12878 cell line [61] for all simulations. The only exception is the simulations for comparing with HiC experiments [20, 47], in which we used the IMR90 cell line instead to match closer to the fibroblast cells in experiments. We labelled beads as HC if the corresponding genomic region contained a peak in the H3K9me3 and/or the LAD data; other beads were marked as EC (see Fig. **1**Aii).

The nuclear lamina (NL) can be modelled in at least two different ways. As conducted in Ref. [62], one could represent the lamina by a smooth, attractive wall using a Lennard-Jones (LJ) potential; however, such a representation can only capture a coarse-grained interaction with the lamina. An alternative approach, implemented here, is to consider the lamina as a layer of beads (*q* = 3), which represents the lamins and other lamina-associated proteins that are part of the NL. This approach can more accurately account for the one-to-one interaction between NL proteins and chromatin. To generate the NL, we put 2500 lamina beads randomly within a 50 nm thick region just beneath the top of the simulation box (Fig. **1**Aiii,B). We chose the lamina beads to be static in the simulation, as it is reasonable to believe that the dynamics of the NL constituents are much slower than that of chromatin.

To probe the structure of the chromosome, we performed molecular dynamics simulations with an implicit solvent (i.e. the nucleoplasm) using a scheme known as Brownian or Langevin dynamics. We simulated the chromosome in a cubic box with a linear dimension *L* = 40*σ* ∼ 2 *μ*m. This size gives a volume fraction of chromatin of about 5%. We employed periodic boundaries in the *x* and *y* direction but fixed boundaries in the *z* direction due to the lamina wall. We used potentials common in polymer physics to simulate the chromatin fibre. First, a purely repulsive Week-Chandler-Andersen (WCA) potential was used to model steric interactions between beads

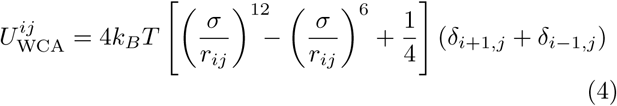

if *r_ij_* < 2^1/6^*σ* and 0 otherwise, where *r_ij_* is the separation between beads *i* and *j*. Second, a finite extensible non-linear elastic (FENE) spring acting between consecutive beads was used to enforce chain connectivity

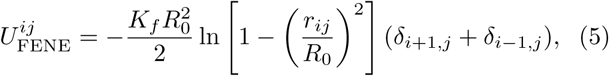

where *R*_0_ = 1.6*σ* is the maximum separation between the beads and *K_f_* = 30*k_B_T*/*σ*^2^ is the spring constant. The superposition of the WCA and FENE potential with the chosen parameters gives a bond length which is approximately equal to *σ* [50]. Third, a Kartky-Porod term was used to model the stiffness of the chromatin fibre

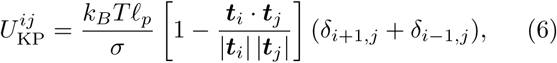

where ***t**_i_* is the tangent vector connecting beads *i* to *i* + 1, and *ℓ_p_* is the persistent length of the chain and is set to 3*σ* ∼ 150 nm, which is within the range of values estimated for chromatin from experiments and computer simulations [63]. The interactions between non-consecutive beads were modelled using a truncated and shifted LJ potential

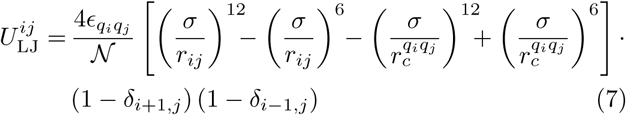

if 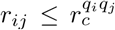 (*q_i_* is the type of bead *i*) and 0 otherwise, where 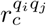 is the cutoff distance of the potential (set to 1.8*σ* for attractive interactions and 2^1/6^*σ* otherwise) and 𝓝 is a normalisation constant to ensure the depth of the potential is equal to the interaction energy *ϵ_q_i_q_j__*. We set *ϵ_q_i_q_j__* = *E*HH (in *k_B_T*) for the interaction between HC beads (*q_i_*, *q_j_* = 2), *ϵ_q_i_q_j__* = *ϵ*_EE_ for that between EC beads (*q_i_*, *q_j_* = 1), and *ϵ_q_i_q_j__* = *ϵ*_HL_ for that between HC and NL beads (*q_i_*, *q_j_* = 2 or 3, *q_i_* ≠ *q_j_*). Other interactions were purely repulsive (*ϵ_q_i_q_j__* = 1). There is no interaction between lamina beads as they are static in the simulation. The sum of potential energy terms involving bead *i* is

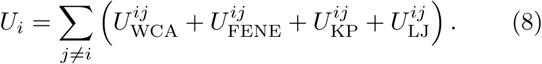

The time evolution of each bead along the fibre is governed by the following Langevin equation

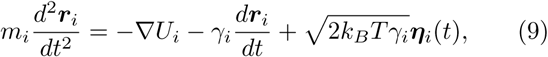

where *m_i_* and *γ_i_* are the mass and the friction coefficient of bead *i*, and ***η**_i_* is its stochastic noise vector with the following mean and variance

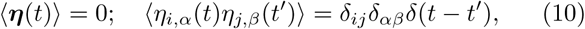

where the Latin and Greek indices run over particles and Cartesian components, respectively. The last term of Eq. (9) represents the random collisions caused by the solvent particles. We assumed all beads have the same mass and friction coefficient (i.e. *m_i_* = *m* and *γ_i_* = *γ*) and set *m* = *γ* = *k_B_T* = 1. We used the Large-scale Atomic/Molecular Massively Parallel Simulator (LAMMPS) (http://lammps.sandia.gov) [64] to numerically integrate the equations of motion using the standard velocity-Verlet algorithm. For the simulation to be efficient yet numerically stable, we set the integration time step to be Δ*t* = 0.01 *τ*_Br_, where *τ*_Br_ is the Brownian time, or the typical time for a bead to diffuse a distance of its own size (i.e. *τ*_Br_ = *σ*^2^/*D* with *D* being the diffusion coefficient).

#### Initial Conditions and Equilibration

We initialised the chromatin fibre as an ideal random walk, in a larger box (*L* = 100*σ*, fixed boundaries) in which the lamina is absent. We allowed the fibre to equilibrate for 10^4^*τ*_Br_, during which the beads can only interact via steric repulsion (with chain connectivity and stiffness maintained). We used the soft potential in the first 4 × 10^3^*τ*_Br_ to remove overlaps in the polymer such that it becomes a self-avoiding chain. In formula, this potential is given by

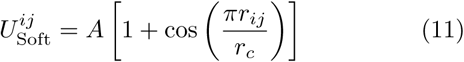

if *r_ij_* < *r_c_* and 0 otherwise, where *r_c_* = 2^1/6^*σ* and *A*, the maximum of the potential, gradually increases from 0 to 20*k_B_T* . We reverted to the WCA potential (for interaction between all beads) for the remaining part of this equilibration period. In the next 5 × 10^3^*τ*_Br_, we compressed the simulation box incrementally to the desired volume (*L* = 40*σ*) using indented walls. We then generated the lamina beads at the top of the simulation box as described above. Finally, we let the chromatin fibre equilibrate with the lamina (interacting via steric repulsion) for 5 × 10^3^*τ*Br, with boundary conditions identical to those for the main simulation run.

### Contact Maps from Simulations and HiC Experiments

The contact maps for the OCI values in Fig. **4** were obtained as follows. In our simulations, we generated HiC-like contact maps by calculating the probability of two beads being in contact, i.e. their separation distance is closer than 5*σ*. This probability was determined from computing the frequency of beads in contact over a 5 × 10^4^*τ*Br time period in each simulation run, which was then averaged over the 20 runs performed for each cell state. For the progeria HiC experiment [47], we used the contact maps (200 kb resolution) directly available from that reference. In particular, we used the Age Control sample for growing and HGPS-p19 sample for progeria. For the senescence experiment [20], we obtained the raw sequencing data for both growing and senescent cells. We used the HiC-Pro pipeline (version 2.10.0) [65] to process the sequencing data and generate the contact maps (50 kb resolution). These maps were normalised using the standard iterative correction procedure [66] to eliminate experimental biases.

### Open Chromatin Index (OCI)

When comparing the contact maps, we considered the open chromatin index (OCI), which is a ratio of the distal contact strength to the local contact strength. More precisely, we defined the (normalised) local contact signal 𝓒_*ℓ*_ and distal contact signal 𝓒_*d*_ for each chromatin bin (say bin *i*) as:

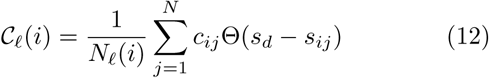

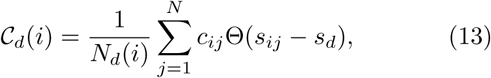

where *c_ij_* is the contact probability between chromatin segments in bins *i* and *j*, *s_ij_* is their genomic separation and *s_d_* = 2 Mb is the genomic distance beyond which we consider contacts to be distal. *N_ℓ_*(*i*) and *N_d_*(*i*) are the number of possible local and distal contact pairs, respectively, for bin *i*. We then used these quantities to define the Open Chromatin Index (OCI) as

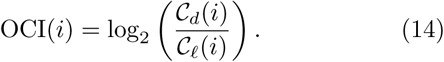

## Acknowledgements

D. Mi., C. B. and D. Ma. acknowledge the European Research Council for funding (Consolidator Grant THREE-DCELLPHYSICS, Ref. 648050). M. C. acknowledges the Carnegie Trust for the Universities of Scotland for PhD studentship funding. T. M. is supported by a Chancellor’s Fellowship held at the University of Edinburgh and the MRC Human Genetics Unit. N. R. was supported by a PhD studentship funded by the Wellcome Trust Sanger Institute and the Royal Thai Government.

## Supplementary Information

### Phase Diagrams

In Fig. **2**, we reported the four possible phases of the system. The figure was constructed by superposing the transition lines calculated from the phase diagrams of the two observables, namely the fraction of beads in contact with the lamina *ψ*(*ϵ*_HH_, *ϵ*_HL_) and the average local number density of beads *ρ*(*ϵ*_HH_, *ϵ*_HL_). These phase diagrams and the corresponding transition lines are shown in Fig. S**1**.

**Figure S 1.**
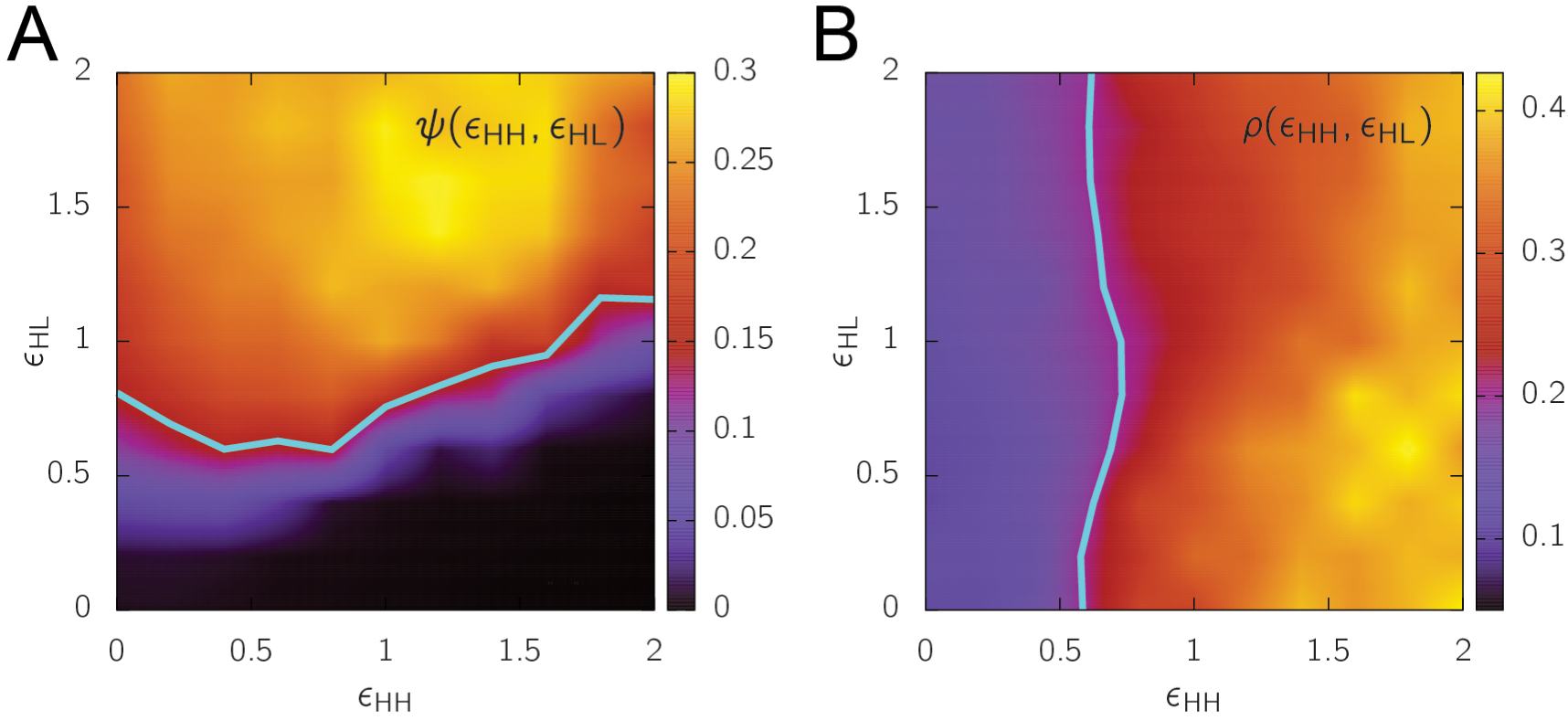
Phases of the simulation model. (A) Measured values for the fraction of beads in contact with the lamina *ψ*(*ϵ*_HH_, *ϵ*_HL_). The transition line (in cyan) was computed based on the cutoff *ψ_c_* = 0.15. (B) Measured values for the average local number density of beads *ρ*(*ϵ*_HH_, *ϵ*_HL_). The transition line was generated based on the cutoff *ρ_c_* = 0.2. These diagrams were created from measuring the observables in the region *ϵ*_HH_, *E*_HL_ ∈ (0, 2) in increments of 0.2 in each direction. At each sampled point, the observables were calculated from averaging over a 5 × 10^4^*τ*_Br_ time period in each simulation run and over 5 runs.

